# Redundancy-selection trade-off in phenotype-structured populations

**DOI:** 10.1101/2021.04.08.439005

**Authors:** Leonardo Miele, R M L Evans, Sandro Azaele

## Abstract

Realistic fitness landscapes generally display a redundancy-fitness trade-off: highly fit trait configurations are inevitably rare, while less fit trait configurations are expected to be more redundant. The resulting sub-optimal patterns in the fitness distribution are typically described by means of effective formulations. However, the extent to which effective formulations are compatible with explicitly redundant landscapes is yet to be understood, as well as the consequences of a potential miss-match. Here we investigate the effects of such trade-off on the evolution of phenotype-structured populations, characterised by continuous quantitative traits. We consider a typical replication-mutation dynamics, and we model redundancy by means of two dimensional landscapes displaying both selective and neutral traits. We show that asymmetries of the landscapes will generate neutral contributions to the marginalised fitness-level description, that cannot be described by effective formulations, nor disentangled by the full trait distribution. Rather, they appear as effective sources, whose magnitude depends on the geometry of the landscape. Our results highlight new important aspects on the nature of sub-optimality. We discuss practical implications for rapidly mutant populations such as pathogens and cancer cells, where the qualitative knowledge of their trait and fitness distributions can drive disease management and intervention policies.

## Introduction

Understanding the interplay between neutrality and selection is considered one of the major challenges in the contemporary theory of biological evolution [1, 2, 3, 4, 5], aiming to bridge the gap between two historically antipodal theories [6]. When neutrality is considered concomitantly with selection, sub-optimal behaviours, that cannot be captured by purely neutralist or selectionist approaches, are expected to emerge due to their interplay [7, 8, 9, 10, 11]. Less fit phenotypes are able to outperform the fittest ones, if they are endowed with higher ‘mutational robustness’ due to some degree of neutrality. This effect is sometimes referred to as the ‘survival-of-the-flattest’ effect, in iconic opposition to the standard ‘survival-of-the-fittest’ paradigm [12, 13]. Although the occurrence of such behaviours is ubiquitous in biology, its characterisation depends crucially both on the genetic architecture and on the mutational topology of the evolving system under investigation [14, 15, 16, 17].

These features have been well documented in the field of molecular phenotype evolution, where the interplay between neutrality and selection is typically described by the redundancy of genotype-phenotype maps [18, 19, 20]. The rate at which mutations occur delineates a major distinction between two possible scenarios, and consequently the kind of mathematical tool suitable for their description. When the mutation rate is low, also known as the ‘weak-mutation’ or monomorphic regime, a complete theory accounting also for neutral effects due to redundancy has been developed in [21].

The complementary, polymorphic, case is generally studied in a deterministic framework. Polymorphic populations are characterised by genetic heterogeneity due to the high mutation rate, so that most of the types are continuously populated (and not the fittest one only). In the polymorphic regime, it is possible to map the low-level genotype dynamics onto the high-level phenotype dynamics only if mutations satisfy a specific condition [22], that is when their rates depend only on the resulting (mutant) phenotype, regardless of the starting (parent) genotype. Although this demanding condition holds for many models of molecular phenotypes, the implications of its violation are much less clear [23].

Phenotype-structured populations belong to the polymorphic category. In such populations, individuals are characterised by (typically) one quantitative trait which is related to reproductive success (fitness) [24]. A common way to model phenotype-structured populations is to describe the quantitative trait of interest by a continuous variable (although discrete versions are possible). Then, mutations are often described by diffusion operators acting on the space of phenotypes. Such properties allow the deterministic mutation-selection dynamics of the population to be described by means of integro-differential equations.

However, diffusion-like mutations do not generally satisfy the special condition [22]; hence, in presence of a degenerate mapping, the two levels of description (phenotypes and fitness) cannot be disentangled and are likely to be different, thus conveying potentially different information about the evolutionary state of the system. In this work, we will study the interplay between neutrality and selection in such rapidly mutating systems.

Phenotypes will be composed of both selective traits (on which fitness depends) and neutral traits (on which it does not), so that the dynamics will be captured by simple fitness landscapes featuring redundancy. Redundancy will be minimally modelled by considering two-dimensional landscapes, where a selective and a neutral trait interact by virtue of a universal redundancy-selection trade-off. Nonetheless, the nature of such trade-offs will be mechanistically different: in the symmetric case, neutrality stems from the property that fitness is given by a combination of the traits composing the phenotype, such combination being degenerate; instead, in the asymmetric case neutrality stems from explicitly considering a completely neutral trait concomitantly with a completely selective trait. Then, redundancy is due to the inherent geometry of the resulting phenotype space, rather than to the degeneracy of the fitness function. For these reasons, we consider the two cases to be suited to qualitatively distinct biological contexts: for instance, the symmetric landscape dates back to the Fisher Geometric Model and has been widely employed in the field of molecular evolution, where the existence of a target optimal configuration of traits is assumed, and any mutation away from it is deleterious [25, 26, 27].

In this work, we will compare phenotype and fitness distributions of populations evolving on both symmetric and asymmetric landscapes. We will derive exact equations governing the resulting fitness dynamics, and compare them to effective formulations. We will show that, despite the fitness distribution on asymmetric landscapes resembling that on symmetric ones, the nature of the two marginal dynamics is crucially different. Particularly, we will demonstrate that in presence of asymmetries between selective and neutral traits, the landscape’s geometry generates contributions that cannot be captured by effective formulations. Finally, we will discuss some biological contexts, where a proper characterisation of neutral contributions to marginal dynamics may be of crucial importance.

## Models and methods

### Redundant fitness landscapes

In molecular evolution, redundancy of genotype-phenotype maps stems from the basic fact that the number of possible genotypes is much larger than that of observed phenotypes, so that such maps must be degenerate. These mappings are also generally strongly biased: some phenotypes are encoded by very few genotypes, whereas most genotypes are organised in networks (that is sets of genotypes connected by a single mutation) that are neutral (i.e. uniformly equally fit), as they map onto the same few phenotypes [28, 29]. It has been argued that this bias should be regarded as a universal feature of any kind of fitness landscapes [30]: ultimately, highly fit individuals are so because they have a phenotype better suited than others to their environment, but such higher functionality will stem from a ‘specific’ (possibly rare) genomic configuration. Hence, a trade-off holds between redundancy and fitness, so that very fit phenotypes would typically not be also highly redundant.

Indeed, in their iconic two-dimensional representation introduced by Wright [31], smooth fitness landscapes exhibit a hill-shaped topography: every phenotype is assigned a height proportional to its fitness, hence the optimum is represented by the top of the hill (see panel **a** of Fig. 1, adapted from [32]). Neutrally related phenotypes, i.e. those sharing the same fitness value, are located at the same height, so that a height contour represents a neutral subset. Since the length of a contour (i.e. the size of the neutral subset) grows with distance from the summit, very fit phenotypes are rare, whereas less fit ones tend to be more abundant. Hence a redundancy-fitness trade-off occurs, akin to that of genotype-phenotype maps.

**Figure 1:**
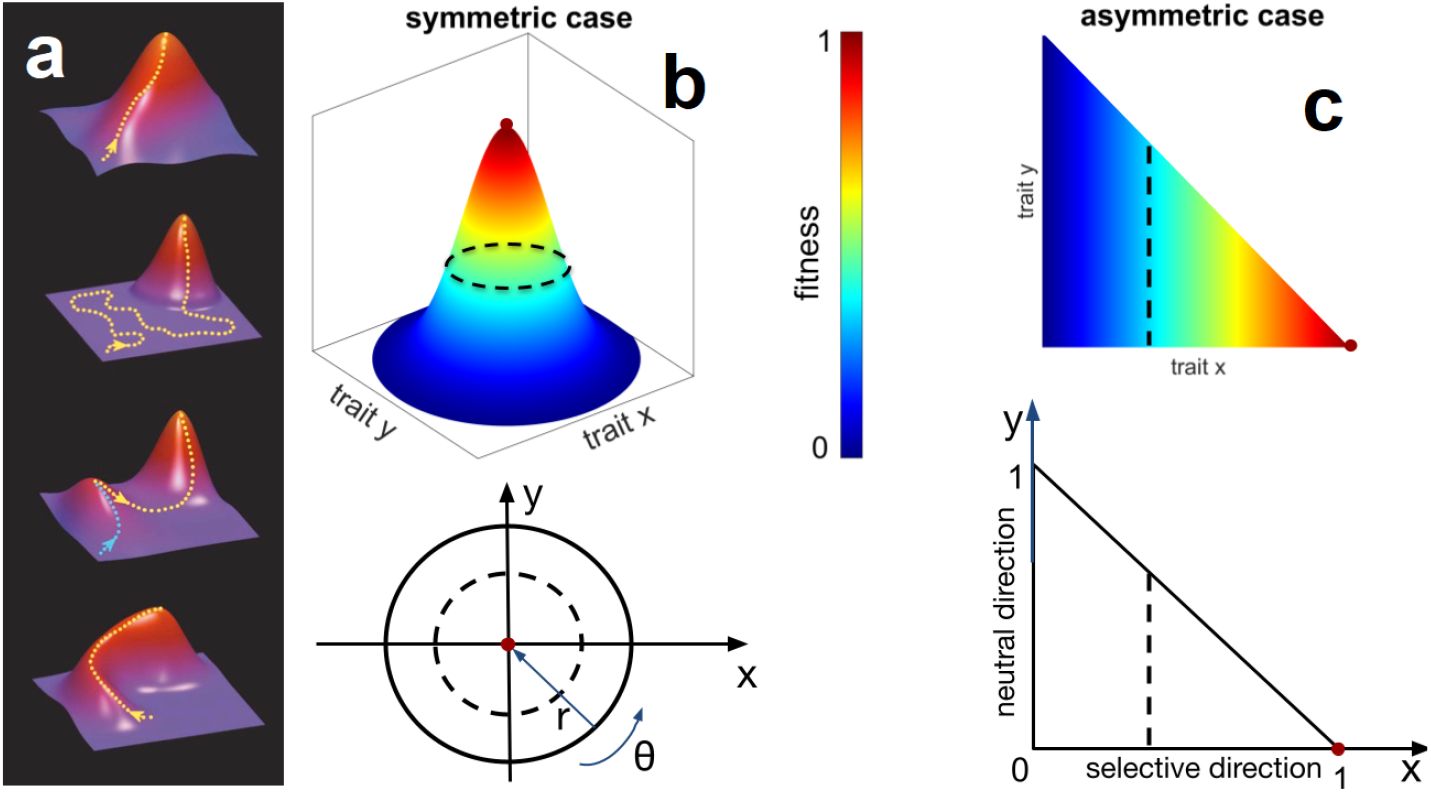
Minimal redundant fitness landscapes. Panel **a**, typical two-dimensional representation of fitness landscapes, exhibiting the redundancy-fitness trade-off: regardless of the topographic details, the size of the neutral subsets decreases as one moves towards the top (adapted from [32]). Panels **b** -**c**: respectively, symmetric and asymmetric redundant fitness landscapes, and projections of the correspondent phenotype spaces, in the trait coordinates (*x, y*). For the symmetric case, fitness depends on the radial distance *r* from the optimum, regardless of the angular position *θ*. For the asymmetric case, fitness is proportional to the trait *x* determining the direction, while the trait *y* is neutral. Dashed black lines represents examples of neutral subsets. Red dots identify the optimum of the respective landscapes. In both cases, the size of the neutral subsets decreases in the selective direction, by virtue of the redundancy-fitness trade-off.

In order to account for the redundancy-fitness trade-off, we shall consider two-dimensional landscapes, but generalisations to higher dimensions are possible. Let *𝒫*_2_ be the phenotype space, and its elements **<u>p</u>** = (*x, y*) ∈ *𝒫*_2_ be the possible phenotypes; the components *x, y* represent respectively the value of the two quantitative traits defining the phenotype. Each phenotype **<u>p</u>** maps into its corresponding fitness value *f* = *F* (**<u>p</u>**) according to the smooth fitness function *F* (**<u>p</u>**); the particular choice of *F* (**<u>p</u>**) determines the fitness landscape of the system. Two phenotypes **<u>p</u>** and **<u>q</u>** are defined to be *neutrally* related if they share the same fitness, that is if *F* (**<u>p</u>**) = *F* (**<u>q</u>**). Then, a *neutral subset* with fitness value *f* is the collection of all neutrally related phenotypes **<u>p</u>** with fitness *F* (**<u>p</u>**) = *f*. For the sake of simplicity we will consider only single-peak landscapes, which have been employed in a variety of biological contexts [33], the study of more complex topographies going beyond the scope of this work.

Redundancy of the landscape is ultimately due to the degeneracy of the fitness function *F*. Here, we shall compare two possible versions of such degeneracy, symmetric (panel **b** Fig. 1) and asymmetric (panel **c** Fig. 1). In panel **b** of Fig. 1, phenotypes are identified by the trait coordinates **<u>p</u>** = (*x, y*). However, their fitness *F* (**<u>p</u>**) depends only on the distance *r*(*x, y*) from the centre. Phenotypes lying on the circle of radius *r* will share the same fitness value regardless of their angular position *θ*, thus forming neutral subsets. Hence, from the pair of trait variables *x* and *y*, we can construct a pair of (respectively) selective and neutral variables (*r, θ*), with which both the phenotype and the fitness dynamics can be described. The phenotype distribution of a population evolving on the symmetric landscape is described by the function *n*(*x, y*) in the original traits coordinates, or equivalently by *n*(*r, θ*) in the corresponding polar coordinates. Given the circular symmetry, the marginal fitness distribution 𝒩^*s*^(*r*) is obtained by integrating the phenotype distribution over the angular coordinate *θ*,

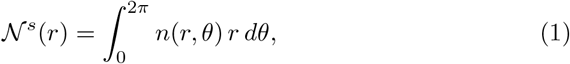

that is the radial distribution. We remark that the landscape exhibits the aforementioned redundancy-fitness trade-off, as the size of neutral subsets varies (linearly in our minimal model) in opposition to fitness.

In the asymmetric case, we assume that the traits *x* and *y* directly express, respectively, selective and neutral effects. So the *x* axis will represent the selective direction, and the *y* axis the neutral direction (panel **c** of Fig. 1), with the fitness function *F* depending on *x* only. The trait space is then closed by the boundary curve *ℬ*(*x*). Neutral subsets are given by vertical lines, that are the collections of points with equal value of the selective trait *x*. From the phenotype distribution *n*(*x, y*) in the original trait coordinates, the marginal fitness distribution *𝒩*^*a*^(*x*) in the asymmetric landscape is given by integration over the neutral variable *y*,

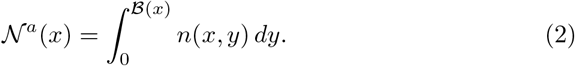

The size of neutral subsets depends on the choice of *ℬ*(*x*): taking a monotonically decreasing function of *x* leads to the desired redundancy-fitness trade-off, equivalent to the symmetric landscape.

### Replicator-Mutator Equation (RME)

The deterministic integro-differential formulation of the mutation-selection dynamics dates back to the ‘continuum-of-alleles’ model introduced by Crow and Kimura [34, 35], and can be derived from stochastic mechanistic models via appropriate continuum limits [24, 36]. Throughout the work, with the generic term ‘individuals’ we refer to the replicating units displaying phenotypic heterogenity, upon which natural selection and mutations act, be they RNA sequences, bacteria or more complex forms of life.

We consider an infinite asexual population. Finite size effects, leading to genetic drift, are thus neglected. The state of the population at time *t* is determined by the phenotype distribution *n*(**<u>p</u>**; *t*). Individuals change their phenotype due to mutation and selection: changes due to mutations are modelled by the Laplacian operator ∇^2^, that is the local diffusion operator acting on the phenotype space *𝒫*_2_, with mutation coefficient *µ*; concomitantly, changes due to selection occur at rate *γ*, and are modelled by the usual replicator term popular in Evolutionary Game Theory [37]. The deterministic temporal evolution of the phenotype distribution *n*(**<u>p</u>**; *t*) for a large population is given by the Replicator-Mutator Equation (RME henceforth):

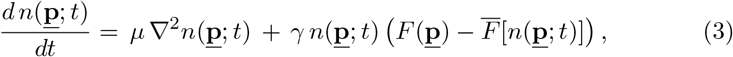

subject to the conditions,

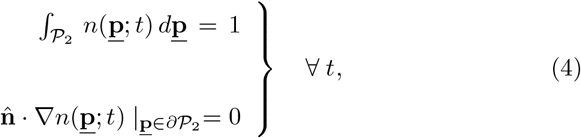

and with 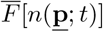 denoting the average fitness of the population at time *t*:

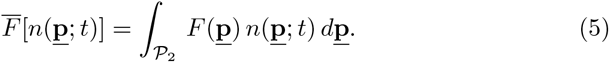

The conditions 4 correspond to the two physical constraints satisfied by the system: conservation of the total population at every time, because neither mutations nor competition alter the number of individuals; and zero flux across the boundaries of the phenotype space, due to reflecting nature of mutations close to the boundary (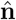 being the unit vector normal to the boundary *∂𝒫*_2_).

The mathematical conditions for which the RME has stationary solutions have been extensively studied [38, 39]. However, explicit analytical solutions are rare because they are hard to obtain (see e.g. [40, 41, 42]). Moreover, multidimensional cases have generally been treated numerically [43]. In order to find the stationary solutions, we employ a self-consistent technique (detailed in the Supplementary Information, section A) that has been applied in similar contexts [25, 44, 45].

Note that, although Eq. 3 contains the timescale *γ*^−1^ and the diffusive coefficient *µ*, the stationary solution will depend on only one relevant parameter 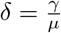, that determines the relative importance of selection and mutation. In the following, we will make simplifying assumptions for the space *𝒫*_2_ and the fitness function *F* (**<u>p</u>**), in order to facilitate analytical calculations on the model. This will allow us to derive useful forms for both the phenotype and the marginal fitness distributions, and compare the differences between symmetric and asymmetric landscapes.

### Simulations

All the analytical results are confirmed by simulating the corresponding finite size stochastic agent-based dynamics. As expected, consistency with the deterministic description is obtained when the population size is very large (order of 10^5^ individuals). The study of finite size effects is possible [46], although it goes beyond the scope of the paper. Simulations have been performed with Java-based language “Processing”, and detailed information can be found in the Supplementary Material, section E. The Processing codes are freely available here.

## Results

### Trait distribution on non-redundant landscapes

Let us first consider a simple one-dimensional case where the fitness landscape is not redundant. This case will provide the baseline results for comparison with the dynamics on redundant landscapes, to elucidate the effects of the redundancy-fitness trade-off.

Let the variable *x* ∈ *𝒫*_1_ = [0, 1] be the single quantitative trait of interest. Let *F* (*x*) be a non-degenerate monotonically increasing function, such that *x* = 1 represents the optimal trait, while *x* = 0 the least fit one. Clearly, since *F* (*x*) is not degenerate, the corresponding fitness landscape is not redundant; each phenotype *x* is uniquely determined by its fitness value. For the sake of simplicity, we shall consider the linear fitness function *F* (*x*) = *x*, for which analytical stationary solutions can be found (mathematical details in the Supplementary Information). However, any monotonic fitness function will produce qualitatively equivalent distributions.

In Fig. 2, we plot the analytical distribution *n*(*x*) for different values of *δ* (solid lines), and compare it with results from numerical simulation (circles and squares). For *δ* = 0, that is in the *purely neutral* scenario, the distribution is flat since every phenotype is equally likely to survive competition, regardless of their fitness value. For *δ >* 0, the distribution is monotonic, always showing an absolute maximum at *x* = 1 (the optimal phenotype), as well as an absolute minimum at *x* = 0 (the least fit one). On increasing *δ* (that is, increasing selection strength or decreasing mutation coefficient), the distribution becomes narrower around the maximum. These profiles represent qualitatively the prediction of the standard *survival-of-the-fittest* paradigm: the most successful phenotype is always the one with the fittest trait, and the population is distributed around the peak of the landscape.

**Figure 2:**
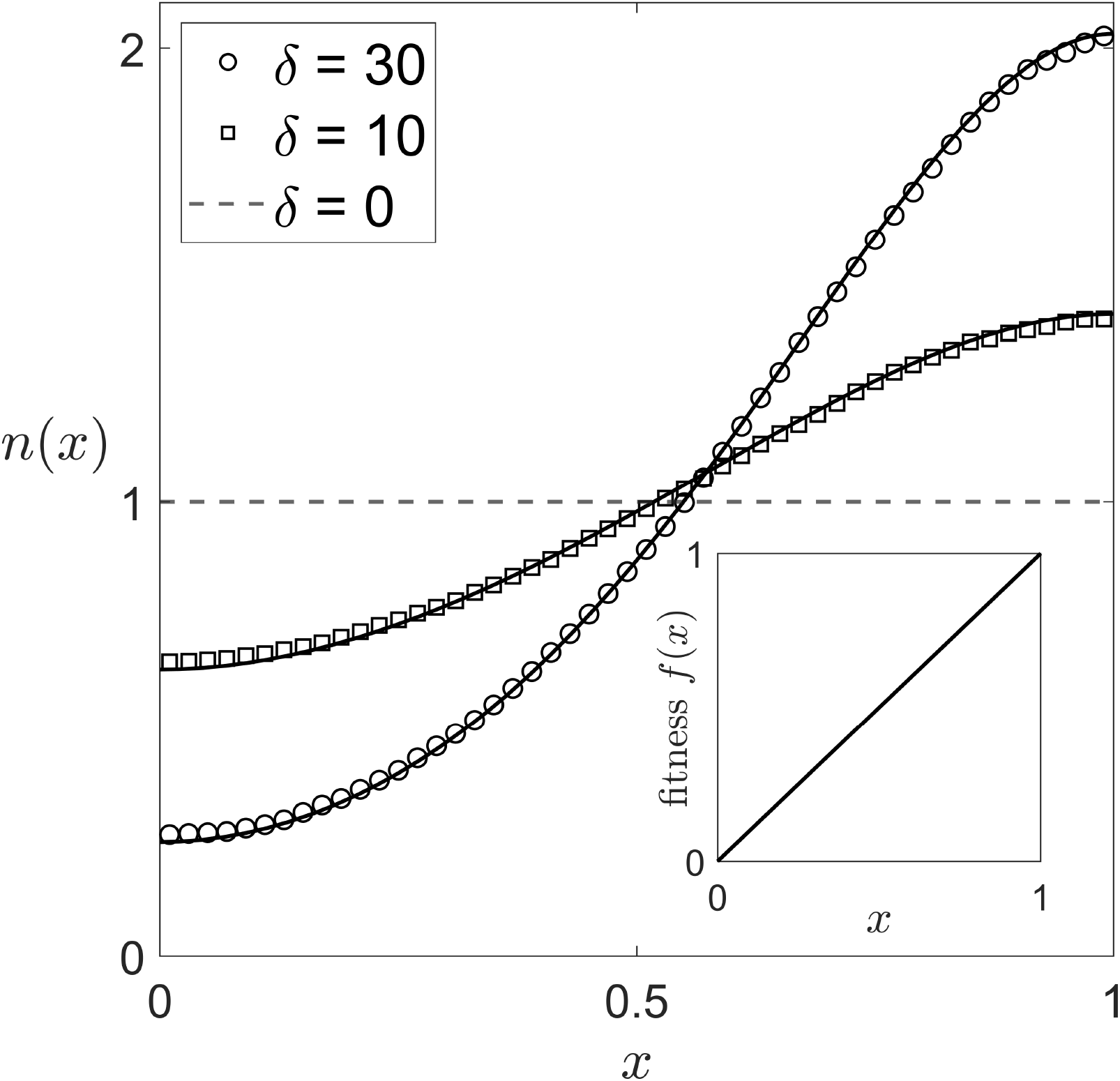
Stationary phenotype distribution on non redundant landscape. Solid lines refer to the analytical solution of the one-dimensional RME, while circles and squares correspond to agent-based numerical simulation of *N* = 10^5^ individuals. With the exception of the neutral case *δ* = 0 (dashed line), the distribution is always monotonically increasing towards the optimal trait *x* = 1, indicating the standard survival-of-the-fittest scenario. Inset: simple fitness landscape for the sole selective variable *x*.

### Trait distribution on redundant landscapes

In redundant landscapes, the phenotype distribution *n*(*x, y*; *t*) evolves in time according to the two-dimensional RME. In general, it is not possible to find an exact closed solution for the stationary distribution. However, in some cases it is possible to obtain spectral solutions. In the following, we shall consider an asymmetric landscape with triangular shape, that is for ℬ(*x*) = 1 − *x* (with 0 ≤ *x* ≤ 1). This specific choice is made in order to facilitate the mathematical tractability of the asymmetric problem. This choice also facilitates the comparison with the symmetric landscape, since the redundancy-selection trade-off decreases linearly with fitness in both cases (see Supplementary Material, sections C and D for mathematical details). However, the same qualitative results are expected to hold for any choice of monotonically decreasing boundary *ℬ* (*x*).

In Fig. 3, we explore the differences between the phenotype distributions *n*(*x, y*) and the marginal fitness distributions *𝒩*^*a,s*^(*f*), at stationarity. The former describes the full distribution of traits over the two-dimensional space *𝒫*_2_. By contrast, the latter describes the one-dimensional distribution of fitness values *f*, and is obtained by integrating the former over the neutral variables.

**Figure 3:**
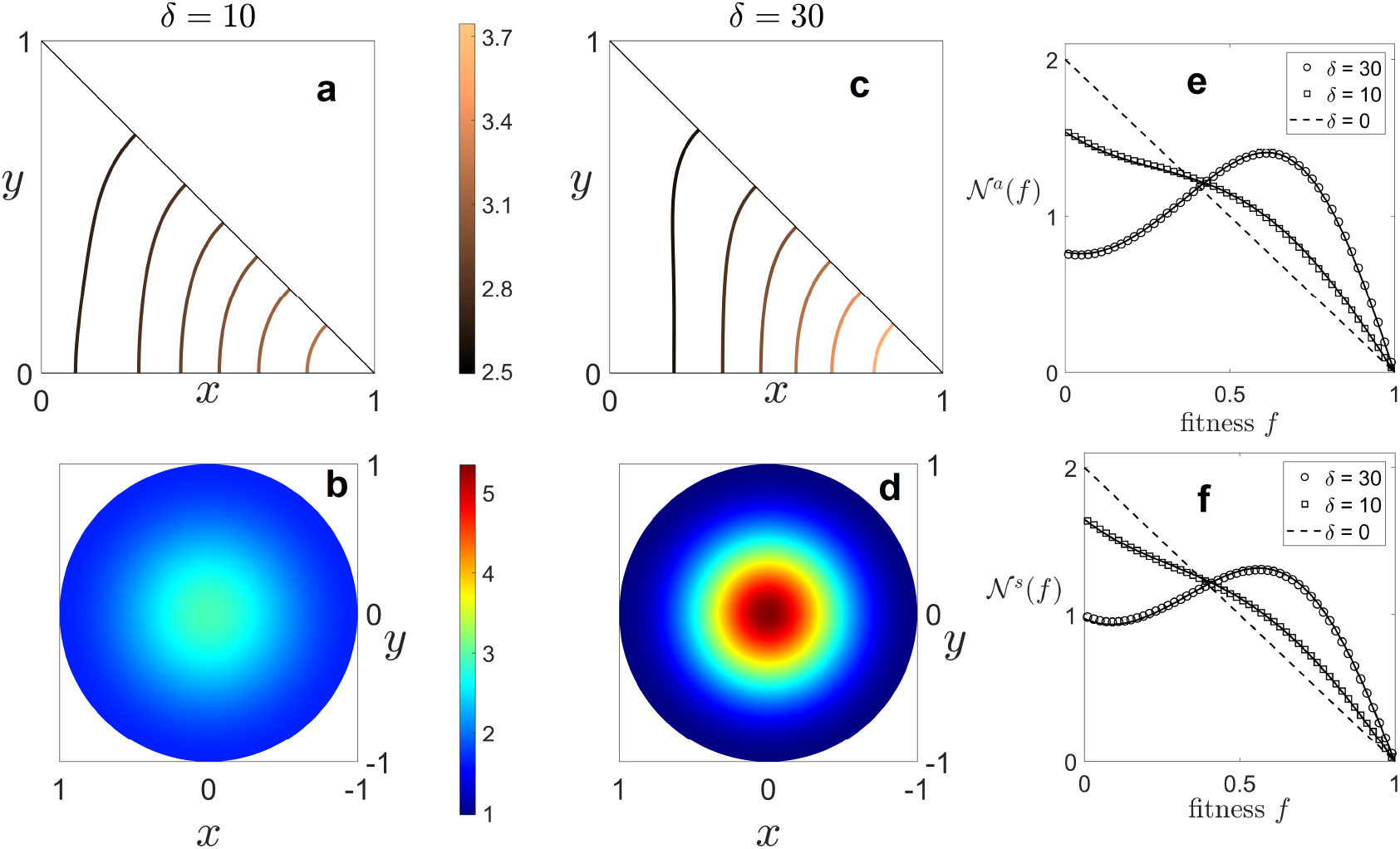
Stationary phenotype distributions and marginal fitness distributions for redundant landscapes. Phenotype distributions: contour lines of iso-density are shown for the asymmetric case (**a** and **c**), while colormaps are shown for the symmetric case (**b** and **d**). In both cases and for every value of *δ >* 0, the distribution has maximum density in correspondence of the optimal trait (that with max fitness), exhibiting a survival-of-the-fittest behaviour. However, the corresponding marginal fitness distributions (**e**-**f**) display rather different behaviours depending on the value of *δ*. Particularly, we distinguish the redundancy-dominated profile (squares *δ* = 10), where the most redundant fitness values are favoured; and the sub-optimal profile (circles *δ* = 30), where the fitness distributions exhibit maximum at an value, smaller than the optimal one. Solid lines refer to analytical solutions of the RME, while scatter plots to agent-based simulations with *N* = 10^5^ individuals.

In panels **a**-**d** of Fig. 3, we plot the analytically obtained phenotype distributions on the trait plane (*x, y*): for the asymmetric case, the iso-density contour lines (**a** and **c**); for the symmetric case, the color-map projection (**b** and **d**). Color code represents the density of *n*(*x, y*), according to the respective color-bars. With the exception of the purely neutral case *δ* = 0, for which the distribution is trivially flat (not shown), the phenotype distributions increase monotonically in the selective direction, i.e. the *x* direction for the asymmetric case, and the radial direction for the symmetric one. In all cases, the distributions display an absolute maximum located at the phenotype with the optimal trait. Similarly to the one-dimensional model, these results again indicate a *survival-of-the-fittest* paradigm, where fitter individuals are more abundant in the population, and the other types are distributed around the optimal with a steepness that increases as *δ* increases.

Let us now consider the behaviour of the marginal fitness distribution *𝒩*^*s*^(*f*) and *𝒩* ^*a*^(*f*) for, respectively, symmetric and asymmetric landscapes. In panels **e**-**f** of Fig. 3, we compare analytical (solid lines) and numerical (circles and squares) profiles of the stationary marginal fitness distributions, for the same values used in the one-dimensional model *δ* = 0, 10, 30.

For *δ* = 0, the purely neutral case, the flat uniform distribution in the two-dimensional phenotype space results in the monotonically decreasing linear profile. Hence, for *δ* = 0 the absolute maximum is found at *x* = 0, which is the most redundant fitness value. Thus, in the absence of selection pressure, fitness values belonging to larger neutral subsets are rewarded, and a scenario consistent with the *survival-of-the-flattest* effect is obtained [47].

For small values of *δ*, the profiles are still monotonically decreasing yet considerably different from the purely neutral case, displaying an increase in the density for intermediate fitness values (see *δ* = 10 case).

For larger values of *δ*, the fitness profile becomes non-monotonic; the previously absolute maximum is now a local one, with the emergence of a new local minimum and of a new absolute maximum. This new absolute peak is located at an intermediate fitness value (see *δ* = 30 case).

In Fig. 4, the positions of the extrema of the fitness profile are shown for a wide range of effective selection pressure values, for asymmetric landscape (the symmetric case is not shown, as it provides the same qualitative result). For *δ* ≲ 14, the profiles are all monotonically decreasing and have an absolute maximum at *f* = 0; we call this regime *redundancy-dominated*, because the most redundant trait is the most abundant in the population. When *δ* crosses a threshold value *δ*_th_, monotonicity is broken, with the emergence of a new peak, that then becomes the absolute maximum at higher *δ*; we call this the *sub-optimal* regime, since the new maximum is located at an intermediate fitness value instead of the optimal one. Increasing selection pressure, the maximum approaches the optimal value *f* = 1, recovering the *survival-of-the-fittest* scenario in the limit of infinite selection pressure. For small values of *δ* in the asymmetric case with linear fitness and triangular shape, a closed analytical approximation of the marginal fitness distribution *𝒩* ^*a*^(*f*) can be obtained. In the Supplementary material, section C, we show that performing a linear perturbation expansion on *δ*, we get:

**Figure 4:**
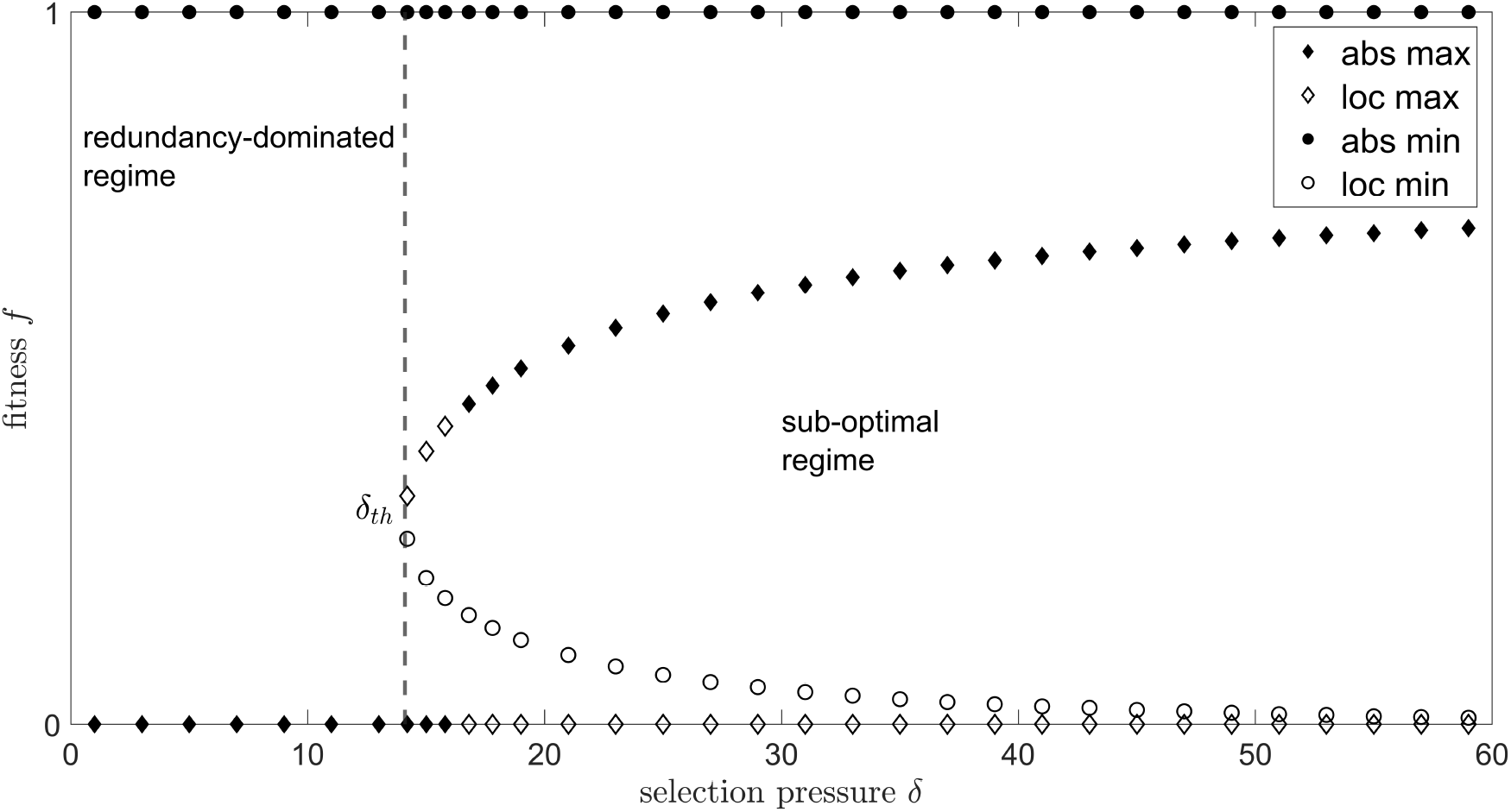
Marginal fitness behaviour. The different regimes of the marginal fitness distribution *𝒩* (*f*) are identified by tracking the extrema of its spectral solution at the variation of selective pressure *δ*. Diamonds (circles) refer to maxima (minima). Filled (empty) symbols refer to absolute (local) extrema. A threshold value *δ*_th_ ≃ 14, estimated with the perturbative solution, separates the two qualitative behaviours. Below *δ*_th_, the fitness distribution is dominated by the most redundant fitness value (redundancy-dominated regime). Above *δ*_th_, the distributions exhibit sub-optimality, as they are dominated by intermediate fitness values. Then, the survival-of-the-fittest scenario is recovered in the limit of very large selection (*δ* → *∞*).

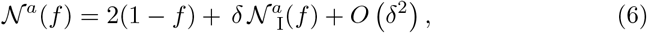

with

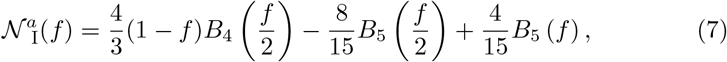

where *B*_*k*_(*z*) is the *k*^th^ Bernoulli polynomial of the variable *z*. This approximation then predicts that the average fitness of the population *ϕ* at stationarity increases linearly with selection pressure, according to:

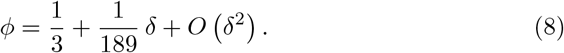

This approximation also predicts the emergence of intermediate local maxima and minima in the marginal fitness distribution for *δ*_th_ ≃ 14 (see Supplementary Figure 1), which is consistent with the results obtained with the spectral solution.

### Marginal fitness dynamics

For the symmetric landscape, the marginal fitness distribution *𝒩* ^*s*^(*f*) is obtained performing the temporal derivative of Eq. 1, and replacing the correspondent RME (details in the Supplementary Information, section D). We find (recall that *f* = 1 − *r*):

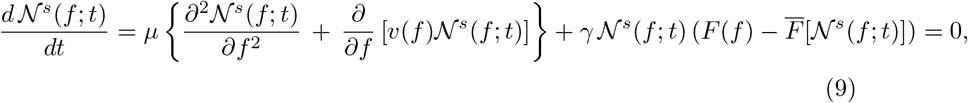

with

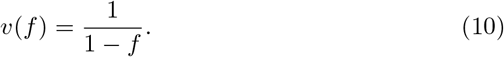

For an asymmetric landscape of general boundary *ℬ* (*x*), the marginal fitness distribution *𝒩*^*a*^(*f* ; *t*) is obtained performing the temporal derivative of Eq. 2, and replacing the correspondent RME (details in the Supplementary Information, section C). In this case, we obtain (recall that *f* = *x*):

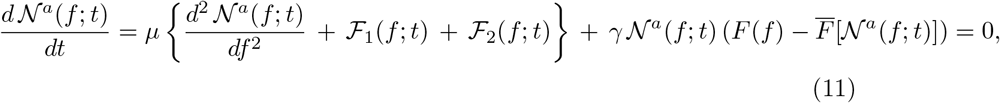

with

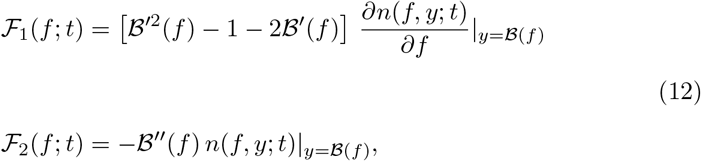

where the prime notation indicates the derivative with respect to the selective variable *f*. The dynamics of the marginal fitness distribution in the symmetric (Eq. 9) and asymmetric (Eq. 11) landscape, display significant differences, which are discussed in detail below.

## Discussion

In this work, we have considered both symmetric (Fig. 1, panel **b**) and asymmetric (Fig. 1, panel **c**) fitness landscapes. Both cases display selective degrees of freedom (namely *x* and *r*), and neutral degrees of freedom (namely *y* and *θ*), which are entwined by a general redundacy-fitness trade-off. However, the different nature of the trade-off generates differences, that are detectable at the marginal fitness dynamics level. Here we shall discuss the consequent analogies and differences, as well as their practical implications.

Contrary to their non redundant counterpart (Fig. 2), we have shown that redundant landscapes display a dual behaviour, depending on the dynamics’ level of description: full phenotype distributions exhibit *survival-of-the-fittest* patterns (Fig. 3, panels **a**-**d**), where most of the population lies in proximity of the landscape optimum; on the other hand, their correspondent marginal fitness distributions may exhibit *sub-optimal* patterns (Fig. 3, panels **e**-**f**), where most of the population displays less fit but more redundant traits (Fig. 4).

For triangular geometry, we have calculated the marginal fitness distribution (Eq. 6) and the average fitness value (Eq. 7), in the weak selection approximation. We observe that the above formulae provide a good estimate of the state of the system up to *δ ≃* 30, above which they break down due to second order selective effects (for details, see Supplementary Material, section C and Supplementary Figure 2). This approximation might also be used as a baseline result to measure landscape’s geometric deviations from the triangular shape.

Acknowledging this duality of behaviours, can help improving the fields in evolutionary epidemiology [48, 49] and cancer dynamics [50, 51], where pathogens are modelled as phenotype-structured populations, and the information on the state of the distributions can be used to design treatment policies.

For example, in a viral or bacterial population, suppose that *x* quantifies the resistance to a drug or antibiotic, so that larger *x* confers higher fitness to its carriers [52]. Then, one might expect the population to be dominated by individuals with highest resistance (i.e. optimal fitness), and a therapy would be developed to counter ‘survival-of-the-fittest’ distributions, hence maximising the intervention on the traits carrying the maximal resistance value. However, if such a selective trait is entwined with another, neutral one (i.e. not affecting the resistance) via a redundancy-fitness trade-off, then the distribution will very likely be dominated by individuals with sub-optimal resistance, and the therapy would erroneously target non-redundant traits, with the possibility of unwittingly helping sub-optimal strains to mutate and become fitter.

On the other hand, suppose that an experimentalist measures the growth rates in a rapidly mutant population as a function of *x*, and obtains a profile similar to panels **e**-**f** of Fig. 3, with a peak in the distribution at an intermediate value 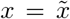 with 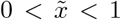. Then they might erroneously conclude that 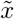 confers the optimal fitness value, whereas, in fact, the trait 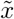 dominates the population due to its redundancy, rather than due to a selective advantage. In the ‘worst case’, by confusing a redundancy-dominated fitness profile with a one-dimensional survival-of-the-fittest distribution, one would infer a direction of selection opposite to the true one, and conclude that trait *x* = 0 has optimal fitness.

In light of the above practical examples, a proper characterisation of neutral contributions is crucial to understand the dual behaviour between full and marginal trait distributions. Neutral information featuring redundant landscapes is often modelled with an effective ‘mutational robustness’ term, where the redundancy-selection trade-off is implicitly accounted for, by introducing some bias to mutations [10, 16, 53, 54, 55]. In these effective formulations, the marginal fitness distribution *𝒩*(*f*) would be governed by some effective RME dynamics depending only on the selective variable *f*, such as

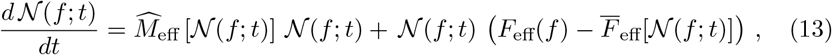

where the interplay between neutrality and selection would be described by either/both a modified ‘mutational operator’ 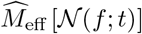, and/or a modified ‘effective fitness’ function *F*_eff_(f) (which is also similar to the case of slowly mutant populations). However, the above effective formulation is not general, and is not appropriate unless the landscape is symmetric.

In this work we have derived the marginal fitness dynamics, by explicit integration over the landscape’s neutral degrees of freedom. In the symmetric landscape, marginalisation leads to a new drift term 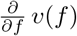, where *v*(*f*) plays the role of a velocity field pushing individuals away from the optimum. This contribution is referred as a ‘mutational entropy’ biasing mutations due to redundancy of the landscape [25, 27]. Thus, the marginal dynamics Eq. 9 is consistent with the effective RME formulation Eq. 13, with:

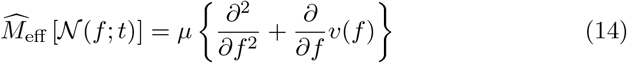

being the new effective mutational operator.

However, in asymmetric landscapes with generic boundary profile *ℬ*(*x*), marginalisation generates contributions of different nature. In Eq. 11, mutations and competition are still captured by, respectively, a local diffusion term and a replicator term. However, marginalisation generates the new contributions *ℱ*_1_(*f* ; *t*) and *ℱ*_2_(*f* ; *t*). The magnitude of such terms depends on the landscape’s geometry, that is on the slope *ℬ*^′^ (*f*) and curvature *ℬ*^″^ (*f*) of the boundary profile. Moreover, from Eq. 12 we observe that these contributions depend on the full phenotype distribution *n*(*f, y*; *t*), thus making the marginal dynamics Eq. 11 an inohomogeneous differential equation. Indeed, the effective formulation Eq. 13 relies on homogeneous differential equations, and it cannot be equivalent to the inhomogeneous one Eq. 11 derived by marginalisation. Therefore, neutral contributions deriving from asymmetric landscapes cannot be identified as ‘effective operators’ acting on the fitness level of description.

This imposes severe limitations on the utility and exactness of effective formulations, for phenotype-structured populations. Indeed, our calculations have shown that solving the high-level fitness dynamics still requires the knowledge of the underlying low-level trait details, and that this issue will occur whenever asymmetries in the trait-space are present.

The new terms due to asymmetry, *ℱ*_1_(*x*; *t*) and *ℱ*_2_(*x*; *t*), have the appearance of effective source contributions to the dynamics, analogous to a spontaneous generation of individuals, if interpreted in the context of a lower-dimensional (non-redundant) fitness landscape. Note that the marginal one-dimensional profiles, shown in Fig. 3 panels **e**-**f**, display a non-zero gradient at the boundaries of the fitness domain, which would require a flux to be present in a truly one-dimensional system. This feature cannot be present in profiles generated by one-dimensional RME models, due to the physical constraints (as, we recall, the total population size is conserved and the system has no flux boundary conditions), unless they are introduced *ad hoc*. We call these emerging sources *effective* because they are generated by the asymmetry in the neutral degrees of freedom, that are unobserved at the marginalised fitness level.

## Conclusions

In this work, we have investigated the RME dynamics of phenotype-structured populations, on minimally redundant landscapes. This kind of dynamics is widely employed in many biological (and other) research areas: population genetics [56], pathogenic evolution [52, 57, 58], RNA evolution [25], game theory [42, 59], language evolution [60]. Its application depends on the identification of rapidly mutating quantitative traits, responsible for phenotypic heterogeneity in the individuals composing the population. Examples of such traits are cytotoxic-drug resistance [61], pathogenic virulence [52, 58] and transmission, antigenic types [62, 63] and hosts’ resistance to infection [64].

Concomitantly with such potential selective traits, accounting for neutral traits is expected to result into asymmetric fitness landscapes, featuring redundancy-selection trade-offs. Particularly, asymmetric landscapes are expected to be found whenever metabolic trade-offs occurs between traits. For instance, the MacArthur’s consumer-resource model [65], is employed to investigate the co-existence of communities competing for a common pool of resources [66, 67]. When multiple resource types are present, the different rates of consumption can be modelled as mutating quantitative traits. If an energetic constraint limits cells’ ability of consumption due to metabolic trade-offs, then the population will evolve on a asymmetric trait space [68].

Similar mechanisms are expected to lead to asymmetric landscapes, in presence of life-history trade-offs. An ideal pathogen would be characterised by high infection transmission, and low induced mortality. In practice, such *super-pathogens* are rarely observed, whereas milder strains are more frequent. This observation is generally explained by acknowledging the existence of a life-history trade-off between transmission and virulence [69], that, in fitness terms, might relate to trade-offs akin to the redundancy-selection one.

Asymmetric landscapes also emerge whenever the phenotype space effectively available is bounded by Pareto-like fronts, outside of which lie all those phenotypic configurations that long-term evolution has excluded, due to their systematic inefficiency [70, 71]. Such trait-spaces have been proposed to explain observed patterns in gene regulation [72], and bacterial growth [73]. Triangular-shaped landscapes, that herein have been used to facilitate calculations, have actually been observed in animal morphology [74, 75, 76]. In game theory, triangular geometries also characterise three-strategies games [77], and have been recently observed to emerge in a numerical study of a rapidly mutant version of the Ultimatum Game [78]. Ultimately, the experimental quantification of the landscape’s asymmetries in the neutral directions is as important as that of selective traits.

In our theoretical work, selection has been introduced by explicitly considering a fitness landscape *F*, and an arbitrary competition rate *γ*. However, in applied contexts, the fitness landscape emerges from the mechanistic interactions associated with the quantitative trait under analysis, whose measurable parameters combine to form effective competition rates [79, 80]. On the other hand, mutations have been modelled by local diffusion over the trait space, characterised by a diffusion coefficient *µ*. Mutations are intended as a global, effective representation of genetic (or higher level) changes that induce phenotypic modification, ignoring the extensive knowledge of the underlying molecular details [81]. This term is appropriate when mutations induce small perturbations on the quantitative traits, i.e. when the components mutate into ‘phenotypically close’ variants. This is not necessarily the case; for instance, when mutations induce a major disruption of the original phenotype, they cannot be modelled by a local diffusion operator (as is the case in the house-of-cards model [82]).

To conclude, we consider our qualitative results to be general and to be relevant whenever rapidly mutant populations evolve on asymmetric redundant fitness landscape. They do not depend on the specifics of the model (which here have been chosen in order to facilitate the mathematical analysis). Our results convey an important message: in general, neutral effects will not be properly captured by effective formulations of mutational robustness; rather, they will generate effective sources at the marginalised fitness-level description. In general, these new contributions will depend on the geometry of the landscape, and the phenotype composition of the population, so that all the microscopic trait information (even for the neutral traits) must be retained in order to properly derive the observable fitness dynamics.

The mathematical procedure herein presented allows the explicit calculation of the trait distribution at stationarity and could be employed to straightforwardly implement redundancy in previous one-dimensional models, so as to include neutral effects. Moreover, it could improve the accuracy of models in evolutionary epidemiology, and the consequent predictions in terms of disease management. As a result, the most effective interventions might not be those that focus on the extremes of the sole fitness-related traits. To interpret such a study, it will be important to consider the relationship between the relevant selective components of traits, as well as their the degree of redundancy in all of the other, neutral, components.

## Acknowledgments

LM is grateful to Robert West and Fabio Peruzzo for fruitful discussion and comments on an earlier version of the manuscript, to Lorenzo Metilli for help with figures, and to Gabriele Lobbia and Giovanni Soldà for computational support. LM, RMLE and SA thank Mauro Mobilia for his co-supervision and feedback. LM thanks the NERC SPHERES DTP (NE/L002574/1) for funding his studentship.

## Notes

### Competing Interest Statement

The authors have declared no competing interest.

## References

[1] A. Wagner, Redundant gene functions and natural selection, Journal of evolutionary biology 12 (1) (1999) 1–16.

[2] S. Ciliberti, O. C. Martin, A. Wagner, Innovation and robustness in complex regulatory gene networks, Proceedings of the National Academy of Sciences 104 (34) (2007) 13591–13596.

[3] A. Wagner, Neutralism and selectionism: a network-based reconciliation, Nature Reviews Genetics 9 (12) (2008) 965.

[4] N. Barghi, J. Hermisson, C. Schlötterer, Polygenic adaptation: a unifying framework to understand positive selection, Nature Reviews Genetics (2020) 1–13.

[5] S. Manrubia, J. A. Cuesta, J. Aguirre, S. E. Ahnert, L. Altenberg, A. V. Cano, P. Catal’ san, R. Diaz-Uriarte, S. F. Elena, J. A. García-Martín, et al., From genotypes to organisms: State-of-the-art and perspectives of a cor-nerstone in evolutionary dynamics, arXiv preprint arXiv:2002.00363.

[6] M. Nei, Mutation-driven evolution, OUP Oxford, 2013.

[7] M. A. Huynen, P. F. Stadler, W. Fontana, Smoothness within ruggedness: the role of neutrality in adaptation, Proceedings of the National Academy of Sciences 93 (1) (1996) 397–401.

[8] D. C. Krakauer, J. B. Plotkin, Redundancy, antiredundancy, and the robustness of genomes, Proceedings of the National Academy of Sciences 99 (3) (2002) 1405–1409.

[9] J. Aguirre, E. L’ sazaro, S. C. Manrubia, A trade-off between neutrality and adaptability limits the optimization of viral quasispecies, Journal of theoretical biology 261 (1) (2009) 148–155.

[10] R. E. Beardmore, I. Gudelj, D. A. Lipson, L. D. Hurst, Metabolic trade-offs and the maintenance of the fittest and the flattest, Nature 472 (7343) (2011) 342.

[11] S. Schaper, A. A. Louis, The arrival of the frequent: how bias in genotype-phenotype maps can steer populations to local optima, PloS one 9 (2).

[12] C. O. Wilke, J. L. Wang, C. Ofria, R. E. Lenski, C. Adami, Evolution of digital organisms at high mutation rates leads to survival of the flattest, Nature 412 (6844) (2001) 331.

[13] J. Sardany’ ses, S. F. Elena, R. V. Solé, Simple quasispecies models for the survival-of-the-flattest effect: The role of space, Journal of Theoretical Biology 250 (3) (2008) 560–568.

[14] M. A. Huynen, Exploring phenotype space through neutral evolution, Journal of molecular evolution 43 (3) (1996) 165–169.

[15] E. Van Nimwegen, J. P. Crutchfield, M. Huynen, Neutral evolution of mutational robustness, Proceedings of the National Academy of Sciences 96 (17) (1999) 9716–9720.

[16] J. A. Draghi, T. L. Parsons, G. P. Wagner, J. B. Plotkin, Mutational robustness can facilitate adaptation, Nature 463 (7279) (2010) 353.

[17] J. Aguirre, J. M. Buldu’ s, M. Stich, S. C. Manrubia, Topological structure of the space of phenotypes: the case of rna neutral networks, PloS one 6 (10).

[18] M. Shackleton, R. Shipma, M. Ebner, An investigation of redundant genotype-phenotype mappings and their role in evolutionary search, in: Proceedings of the 2000 Congress on Evolutionary Computation. CEC00 (Cat. No. 00TH8512), Vol. 1, IEEE, 2000, pp. 493–500.

[19] F. M. Codoner, J.-A. Dar’ sos, R. V. Solé, S. F. Elena, The fittest versus the flattest: experimental confirmation of the quasispecies effect with subviral pathogens, PLoS pathogens 2 (12) (2006) e136.

[20] A. Wagner, The origins of evolutionary innovations: a theory of transformative change in living systems, OUP Oxford, 2011.

[21] B. S. Khatri, R. A. Goldstein, A coarse-grained biophysical model of sequence evolution and the population size dependence of the speciation rate, Journal of theoretical biology 378 (2015) 56–64.

[22] K. Sato, K. Kaneko, Evolution equation of phenotype distribution: General formulation and application to error catastrophe, Physical Review E 75 (6) (2007) 061909.

[23] B. Khatri, Survival of the frequent at finite population size and mutation rate: filing the gap between quasispecies and monomorphic regimes doi:10.1101/375147.

[24] R. H. Chisholm, T. Lorenzi, L. Desvillettes, B. D. Hughes, Evolutionary dynamics of phenotype-structured populations: from individual-level mechanisms to population-level consequences, Zeitschrift für angewandte Mathematik und Physik 67 (4) (2016) 100.

[25] L. S. Tsimring, H. Levine, D. A. Kessler, Rna virus evolution via a fitness-space model, Physical review letters 76 (23) (1996) 4440.

[26] H. A. Orr, The distribution of fitness effects among beneficial mutations in fisher’s geometric model of adaptation, Journal of theoretical biology 238 (2) (2006) 279–285.

[27] U. Gerland, T. Hwa, On the selection and evolution of regulatory dna motifs, Journal of molecular evolution 55 (4) (2002) 386–400.

[28] J. M. Smith, Natural selection and the concept of a protein space, Nature 225 (5232) (1970) 563–564.

[29] A. Wagner, The role of robustness in phenotypic adaptation and innovation, Proceedings of the Royal Society B: Biological Sciences 279 (1732) (2012) 1249–1258.

[30] B. S. Khatri, R. A. Goldstein, Biophysics and population size constrains speciation in an evolutionary model of developmental system drift, PLoS computational biology 15 (7) (2019) e1007177.

[31] S. Wright, The roles of mutation, inbreeding, crossbreeding, and selection in evolution, Vol. 1, na, 1932.

[32] F. J. Poelwijk, D. J. Kiviet, D. M. Weinreich, S. J. Tans, Empirical fitness landscapes reveal accessible evolutionary paths, Nature 445 (7126) (2007) 383.

[33] M.-E. Gil, F. Hamel, G. Martin, L. Roques, Dynamics of fitness distributions in the presence of a phenotypic optimum: an integro-differential approach, Nonlinearity 32 (10) (2019) 3485.

[34] J. F. Crow, M. Kimura, The theory of genetic loads, in: Proceedings of the XIth International Congress of Genetics, Vol. 2, Pergamon Press Oxford, 1964, pp. 495–505.

[35] M. Kimura, A stochastic model concerning the maintenance of genetic variability in quantitative characters., Proceedings of the National Academy of Sciences of the United States of America 54 (3) (1965) 731.

[36] N. Champagnat, R. Ferrière, S. M’ seléard, Unifying evolutionary dynamics: from individual stochastic processes to macroscopic models, Theoretical population biology 69 (3) (2006) 297–321.

[37] P. Schuster, K. Sigmund, Replicator dynamics, Journal of theoretical biology 100 (3) (1983) 533–538.

[38] R. Bürger, I. M. Bomze, Stationary distributions under mutation-selection balance: structure and properties, Advances in applied probability 28 (1) (1996) 227–251.

[39] R. Bürger, Mathematical properties of mutation-selection models, Genetica 102 (1998) 279.

[40] M. Alfaro, R. Carles, Replicator-mutator equations with quadratic fitness, Proceedings of the American Mathematical Society 145 (12) (2017) 5315–5327.

[41] M. Alfaro, M. Veruete, Evolutionary branching via replicator–mutator equations, Journal of Dynamics and Differential Equations 31 (4) (2019) 2029–2052.

[42] M. Ruijgrok, T. W. Ruijgrok, An effective replicator equation for games with a continuous strategy set, Dynamic Games and Applications 5 (2) (2015) 157–179.

[43] Y. Cohen, Evolutionary distributions, Evolutionary Ecology Research 11 (4) (2009) 611–635.

[44] I. M. Rouzine, J. Wakeley, J. M. Coffin, The solitary wave of asexual evolution, Proceedings of the National Academy of Sciences 100 (2) (2003) 587–592.

[45] O. Hallatschek, The noisy edge of traveling waves, Proceedings of the National Academy of Sciences 108 (5) (2011) 1783–1787.

[46] A. Ardašseva, A. R. Anderson, R. A. Gatenby, H. M. Byrne, P. K. Maini, T. Lorenzi, Comparative study between discrete and continuum models for the evolution of competing phenotype-structured cell populations in dynamical environments, Physical Review E 102 (4) (2020) 042404.

[47] C. O. Wilke, Quasispecies theory in the context of population genetics, BMC evolutionary biology 5 (1) (2005) 44.

[48] A. P. Galvani, Epidemiology meets evolutionary ecology, Trends in Ecology & Evolution 18 (3) (2003) 132–139.

[49] T. Day, T. Parsons, A. Lambert, S. Gandon, The price equation and evolutionary epidemiology, Philosophical Transactions of the Royal Society B 375 (1797) (2020) 20190357.

[50] R. V. Sol’ se, T. S. Deisboeck, An error catastrophe in cancer?, Journal of Theoretical Biology 228 (1) (2004) 47–54.

[51] J. Clairambault, An evolutionary perspective on cancer, with applications to anticancer drug resistance modelling and perspectives in therapeutic control, J. Math. Study 52 (4) (2019) 470–496.

[52] T. Day, S. R. Proulx, A general theory for the evolutionary dynamics of virulence, The American Naturalist 163 (4) (2004) E40–E63.

[53] D. De Martino, F. Capuani, A. De Martino, Growth against entropy in bacterial metabolism: the phenotypic trade-off behind empirical growth rate distributions in e. coli, Physical biology 13 (3) (2016) 036005.

[54] A. De Martino, T. Gueudr’ se, M. Miotto, Exploration-exploitation tradeoffs dictate the optimal distributions of phenotypes for populations subject to fitness fluctuations, Physical Review E 99 (1) (2019) 012417.

[55] E. Rigato, G. Fusco, Effects of phenotypic robustness on adaptive evolutionary dynamics, Evolutionary Biology 47 (3) (2020) 233–239.

[56] J. Y. Wakano, T. Funaki, S. Yokoyama, Derivation of replicator–mutator equations from a model in population genetics, Japan Journal of Industrial and Applied Mathematics 34 (2) (2017) 473–488.

[57] A. Korobeinikov, C. Dempsey, A continuous phenotype space model of rna virus evolution within a host, Mathematical Biosciences & Engineering 11 (4) (2014) 919.

[58] L. Bolzoni, G. A. De Leo, Unexpected consequences of culling on the eradication of wildlife diseases: the role of virulence evolution, The American Naturalist 181 (3) (2013) 301–313.

[59] I. M. Bomze, R. Burger, Stability by mutation in evolutionary games, Games and Economic Behavior 11 (2) (1995) 146–172.

[60] K. M. Page, M. A. Nowak, Unifying evolutionary dynamics, Journal of theoretical biology 219 (1) (2002) 93–98.

[61] T. Lorenzi, R. H. Chisholm, J. Clairambault, Tracking the evolution of cancer cell populations through the mathematical lens of phenotype-structured equations, Biology direct 11 (1) (2016) 43.

[62] A. Sasaki, Evolution of antigen drift/switching: continuously evading pathogens, Journal of Theoretical Biology 168 (3) (1994) 291–308.

[63] A. Sasaki, Y. Haraguchi, Antigenic drift of viruses within a host: a finite site model with demographic stochasticity, Journal of Molecular Evolution 51 (3) (2000) 245–255.

[64] T. Lorenzi, A. Pugliese, M. Sensi, A. Zardini, Evolutionary dynamics in an si epidemic model with phenotype-structured susceptible compartment, arXiv preprint arXiv:2010.10443.

[65] R. MacArthur, Species packing and competitive equilibrium for many species, Theoretical population biology 1 (1) (1970) 1–11.

[66] L. Pacciani-Mori, A. Giometto, S. Suweis, A. Maritan, Dynamic metabolic adaptation can promote species coexistence in competitive communities, PLoS computational biology 16 (5) (2020) e1007896.

[67] D. Gupta, S. Garlaschi, S. Suweis, S. Azaele, A. Maritan, An effective resource-competition model for species coexistence (2021). arXiv:2104.01256.

[68] M. Amicone, I. Gordo, Molecular signatures of resource competition: clonal interference drives the emergence of ecotypes, bioRxiv.

[69] S. Alizon, A. Hurford, N. Mideo, M. Van Baalen, Virulence evolution and the trade-off hypothesis: history, current state of affairs and the future, Journal of evolutionary biology 22 (2) (2009) 245–259.

[70] O. Shoval, H. Sheftel, G. Shinar, Y. Hart, O. Ramote, A. Mayo, E. Dekel, K. Kavanagh, U. Alon, Evolutionary trade-offs, pareto optimality, and the geometry of phenotype space, Science 336 (6085) (2012) 1157–1160.

[71] B. Xue, P. Sartori, S. Leibler, Environment-to-phenotype mapping and adaptation strategies in varying environments, Proceedings of the National Academy of Sciences 116 (28) (2019) 13847–13855.

[72] A. Y. Weiße, D. A. Oyarzu’ sn, V. Danos, P. S. Swain, Mechanistic links between cellular trade-offs, gene expression, and growth, Proceedings of the National Academy of Sciences 112 (9) (2015) E1038–E1047.

[73] S. Klumpp, T. Hwa, Bacterial growth: global effects on gene expression, growth feedback and proteome partition, Current opinion in biotechnology 28 (2014) 96–102.

[74] G. R. McGhee, The geometry of evolution: adaptive landscapes and theoretical morphospaces, Cambridge University Press, 2006.

[75] E. O. Wilson, Caste and division of labor in leaf-cutter ants (hymenoptera: Formicidae: Atta), Behavioral ecology and sociobiology 7 (2) (1980) 157–165.

[76] U. M. Norberg, J. M. Rayner, Ecological morphology and flight in bats (mammalia; chiroptera): wing adaptations, flight performance, foraging strategy and echolocation, Philosophical Transactions of the Royal Society of London. B, Biological Sciences 316 (1179) (1987) 335–427.

[77] A. Boccabella, R. Natalini, L. Pareschi, On a continuous mixed strategies model for evolutionary game theory, Kinetic & Related Models 4 (1) (2011) 187.

[78] R. Evans, Pay-off scarcity causes evolution of risk-aversion and extreme altruism, Scientific reports 8 (1) (2018) 1–10.

[79] T. Day, S. Gandon, Insights from price’s equation into evolutionary, Disease evolution: models, concepts, and data analyses 71 (2006) 23.

[80] T. Day, S. Gandon, Applying population-genetic models in theoretical evolutionary epidemiology, Ecology Letters 10 (10) (2007) 876–888.

[81] G. Martin, S. F. Elena, T. Lenormand, Distributions of epistasis in microbes fit predictions from a fitness landscape model, Nature genetics 39 (4) (2007) 555–560.

[82] J. F. Kingman, A simple model for the balance between selection and mutation, Journal of Applied Probability 15 (1) (1978) 1–12.

