## Supplementary Material for "Redundancy-selection trade-off in phenotype-structured populations"

### A Self-consisted method to solve stationary RME

We recall that the temporal dynamics of the phenotype-structured population characterised by mutation diffusion coefficient $\mu$ and competition rate $\gamma$, is given by the following Replicator-Mutator Equation (RME):

$$\frac{d\,n(\underline{\mathbf{p}};t)}{dt} = \mu\,\nabla^2 n(\underline{\mathbf{p}};t) + \gamma\,n(\underline{\mathbf{p}};t)\left(F(\underline{\mathbf{p}}) - \overline{F}[n(\underline{\mathbf{p}};t)]\right), \tag{1}$$

where $\underline{\mathbf{p}}$ is the set of quantitative traits determining the phenotype, $n(\underline{\mathbf{p}};t)$ is the density of individuals with phenotype $\mathbf{p}$ at time $t$, $F(\underline{\mathbf{p}})$ is the fitness function, and $\bar{F}$ is the average fitness. When the system has reached stationarity, the population's average fitness $\bar{F}[n(\underline{\mathbf{p}};t)]$ does not change in time. Despite its value being unknown, the nonlinearity of the RME can be managed by treating the average fitness as a parameter that we will call $\phi$. Then, one can look for the *$\phi$-family* of solutions of the linear RME

$$\mu \nabla^2 n_\phi(\underline{\mathbf{p}}) + \gamma\, n_\phi(\underline{\mathbf{p}})\big(F(\underline{\mathbf{p}}) - \phi\big) = 0, \tag{2}$$

subject to the conditions given in Eq. 4 of the main text. Finally, the correct solution can be identified self-consistently by solving the constraint:

$$\phi = \int_{\mathcal{P}_2} F(\underline{\mathbf{p}})\, n_\phi(\underline{\mathbf{p}})\, d\underline{\mathbf{p}}. \tag{3}$$

Such a procedure has a general validity and can be employed whenever a solution of the linear RME 2 is available. Solving the linear Eq. 2 is, in principle, an easier task than the nonlinear version. However, the choice of method will depend on the properties of both the fitness landscape $F$ and the phenotype space $\mathcal{P}_2$, that in turn affect the boundary conditions of the problem.

### B Trait distribution on non-redundant landscapes

Here we show how to obtain the solution for the simple non-redundant landscape $\mathcal{P}_1 = [0, 1]$, with linear fitness function. We recall that the one-dimensional stationary phenotype distribution $n(x)$ with linear fitness function $F(x) = x$, is solution of the following Replicator-Mutator Equation (RME):

$$\mu \frac{\partial^2 n(x)}{\partial x^2} + \gamma\, n(x)\Big(x - \phi\Big) = 0, \tag{4}$$

with:

$$\begin{cases} \int_0^1 n(x)\, dx = 1 \\ \frac{\partial n(x)}{\partial x}\, |_{x=0} = \frac{\partial n(x)}{\partial x}\, |_{x=1} = 0 \\ \phi = \int_0^1 x\, n(x)\, dx. \end{cases} \tag{5}$$

In Eq. 5, the first condition ensures normalisation of the solution; the second corresponds to no-flux boundary conditions due to reflecting mutations at the boundary of the domain; the third condition fixes the average fitness that the solution will have to satisfy self-consistently.

Performing the transformation of variable $z = \sqrt[3]{\frac{\gamma}{\mu}}\,(\phi - x)$, Eq. 4 becomes:

$$\frac{\partial^2 n(z)}{\partial z^2} - z\, n(z) = 0, \tag{6}$$

that is the well known Airy differential equation (Abramowitz and Stegun 1964). The solution is then a linear combination of the Airy functions of first and second kind which, back in the original variable, reads:

$$n(x) = \mathcal{Z}\Big\{ Ai\left[\sqrt[3]{\delta}\,(\phi - x)\right] + \mathcal{C}\, Bi\left[\sqrt[3]{\delta}\,(\phi - x)\right]\Big\}, \quad \delta = \frac{\gamma}{\mu}, \tag{7}$$

with $\mathcal{Z}$ and $\mathcal{C}$ constants of integration given by (prime notation indicates the $x$ derivative):

$$\begin{cases} \mathcal{C} = -\frac{Ai'\left(\sqrt[3]{\delta}\phi\right)}{Bi'\left(\sqrt[3]{\delta}\phi\right)} \\ \mathcal{Z}^{-1} = \int_0^1 Ai\left[\sqrt[3]{\delta}\,(\phi - x)\right] + \mathcal{C}\, Bi\left[\sqrt[3]{\delta}\,(\phi - x)\right]\, dx. \end{cases} \tag{8}$$

Equation 8 represents the self-consistent fitness (and phenotype) distribution at stationarity of the one dimensional RME. In order to plot the equation, the proper average fitness value $\phi$ is computed numerically by using the last condition in Eq. 5, as explained in section A.

### C Trait distribution on asymmetric redundant landscape

#### Spectral solution

In the asymmetric case, the dynamics takes place in the phenotype space $\mathcal{P}_2 = \{0 \le x \le 1, 0 \le y \le \mathcal{B}(x)\}$. The $x$ axis represents the selective direction, while the $y$ axis the neutral one. The fitness landscape is bounded by the profile $y = \mathcal{B}(x)$. The optimal phenotype is the edge $(1, 0)$, and has maximum fitness value $f = 1$. In this case, the stationary phenotype distribution $n(x, y)$ solves the following two dimensional RME:

$$\mu \nabla^2 n(x,y) \; + \; \gamma \, n(x,y)\Big(F(x) - \phi\Big) = 0, \tag{9}$$

subject to conditions:

$$\begin{cases} \int_0^1 \int_0^{1-x} n(x,y)\, dxdy = 1 & \textit{population conservation} \\ \frac{\partial n(x,y)}{\partial x} \,|_{x=0} = \frac{\partial n(x,y)}{\partial y} \,|_{y=0} = \frac{\partial n(x,y)}{\partial x} + \frac{\partial n(x,y)}{\partial y} \,|_{y=1-x} = 0 & \textit{no-flux} \\ \phi = \int_0^1 \int_0^{1-x} F(x)\, n(x,y)\, dxdy & \textit{self-consistent average fitness.} \end{cases} \tag{10}$$

Despite the apparent simple form of Eq. 9, the separation of variables method fails due to the non trivial no-flux condition at the boundary $y = 1 - x$. Thus, we look for a solution of the form:

$$n(x,y) = \sum_{i,j} a_{i,j} e_{i,j}(x,y), \tag{11}$$

where the $e_{i,j}(x, y)$ are the eigenfunctions of the Helmholtz problem on the triangle with Neumann boundary conditions:

$$\begin{cases} \nabla^2 \, e_{i,j}(x,y) = \lambda_{i,j} \, e_{i,j}(x,y) \\ \frac{\partial e_{i,j}(x,y)}{\partial x} \,|_{x=0} = \frac{\partial e_{i,j}(x,y)}{\partial y} \,|_{y=0} = \frac{\partial e_{i,j}(x,y)}{\partial x} + \frac{\partial e_{i,j}(x,y)}{\partial y} \,|_{y=1-x} = 0. \end{cases} \tag{12}$$

The above problem is solved by Prager's method (Damle and Peterson 2010): first, the eigenfunctions corresponding to the same problem on the square domain are found; then these functions are "folded" around the diagonal with a linear transformation. The resulting normalised eigenfunctions and relative eigenvalues are:

$$\begin{cases} e_{0,0} = 2 \\ e_{i,j}(x,y) = \mathcal{Z}_{i,j}\big[\cos(i\pi x)\,\cos(j\pi y)\,+(-1)^{i+j}\,\cos(i\pi y)\,\cos(j\pi x)\big] \quad \forall i,j\neq 0 \\ \lambda_{i,j} = -\pi^2\,(i^2+j^2)\,, \end{cases} \tag{13}$$

with $\mathcal{Z}_{i,j} = \sqrt{2}+\big(2-\sqrt{2}\big)\,\delta_{i,j}+\big(2-\sqrt{2}\big)\,\delta_{j,0}$ constant of normalisation. The above eigenfunctions have the same properties:

$$\begin{cases} \int_0^1\int_0^{1-x} e_{i,j}\,e_{k,l}\,dxdy = \delta_{i,k}\,\delta_{j,l} & \textit{orthonormality} \\ e_{i,j}(x,y) = (-1)^{i+j}\,e_{j,i}(x,y) & \textit{symmetry} \\ \int_0^1\int_0^{1-x} e_{i,j}(x,y)\,dxdy = 0 \quad \forall\, i,j\neq 0. \end{cases} \tag{14}$$

Due to the symmetry property, only half of the $\{e_{i,j}\}$ are needed to create a complete orthonormal basis for $L^2$. Hence, the expansion reads:

$$n(x,y) = \sum_{i=1}^{+\infty}\sum_{j=0}^{i} a_{i,j}e_{i,j}(x,y). \tag{15}$$

Combining the normalisation constraint on the solution with the last property in Eq. 14, we get:

$$\int_0^1\int_0^{1-x} n(x,y)\,dxdy = a_{0,0}+0+\cdots+0 = 1. \tag{16}$$

Hence, the zero coefficient of the expansion is $a_{0,0}=1$. In order to find the remaining coefficients we insert the expansion Eq. 15 into Eq. 9, then we multiply by $e_{k,l}(x,y)$ and we integrate over the triangle domain. Using the orthonormalisation condition and rearranging the terms we get:

$$\big(\lambda_{k,l}-\delta\phi+\delta\langle\,F(x)\,\rangle_{k,l;k,l}\big)\,a_{k,l}\,+\,\delta\sum_{i\neq k}^{+\infty}\sum_{j\neq l}^{i} a_{i,j}\,\langle\,F(x)\,\rangle_{i,j;k,l} = -\delta\,\langle\,F(x)\,\rangle_{0,0;k,l}, \tag{17}$$

where the notation $\langle\,\cdot\,\rangle_{i,j;k,l}$ indicates the average over $e_{i,j}(x,y)\,e_{k,l}(x,y)$. The expression Eq. 17 represents an infinite system of linear equations for the unknown coefficients $\{a_{i,j}\}$ (we remind that at this stage the coefficients still depend on the average fitness $\phi$).

For general values of $\delta$, one needs to truncate the system to a finite number of eigenfunctions, then solve the linear problem with standard numerical routines (such as the MATLAB function *linsolve*). Once the $\phi-$dependant coefficients are found, the solution is obtained by solving numerically the self-consistent condition in Eq. 10 regarding the average fitness.

However, for small values of $\delta$ it is possible to find a closed approximation of the marginal fitness distribution in the case of linear fitness function, as will be explained in the next subsection.

#### Perturbative solution

We note that the solution truncated to the zero$^{\text{th}}$ eigenfunction $n(x,y) = a_{0,0}\, e_{0,0} = 2$ corresponds to the solution of the neutral dynamics, i.e. to the case $\delta = 0$. For small values of $\delta$, i.e. for weak selection, we expect the neutral solution to be perturbed by contributions due to selection. In order to find the first order corrections, we assume that:

$$a_{i,j} = O\left(\delta\right) \quad \forall i, j \neq 0. \tag{18}$$

Inserting this assumption into the system Eq. 17 and retaining only the terms of first order in $\delta$, we get:

$$a_{i,j} = -\delta\, \frac{\langle\, F(x)\,\rangle_{0,0;i,j}}{\lambda_{i,j}}. \tag{19}$$

Computing the values of the averages for the linear fitness function $F(x) = x$, and of the eigenvalues we obtain:

$$\begin{aligned} n(x,y) &= 2 + \delta\Big\{ -4\sum_{k=1}^{+\infty} \frac{1}{k^4\pi^4} e_{k,0}(x,y) + 4\sum_{k=1}^{+\infty} \frac{(-1)^k}{k^4\pi^4} e_{k,k}(x,y)\Big\} + O(\delta^2) \qquad (20) \\ &= 2 + \delta\Big\{ -4\sum_{k=1}^{+\infty} \frac{1}{k^4\pi^4}\left[\cos(k\pi x) + (-1)^k\cos(k\pi y)\right] + 4\sum_{k=1}^{+\infty} \frac{(-1)^k}{k^4\pi^4}\cos(k\pi x)\cos(k\pi y)\Big\} + O(\delta^2). \end{aligned}$$

Integrating Eq. 20 over the neutral variable $y$ and replacing the selective variable $x$ with its fitness value $f$ (recall that $f = x$), we obtain the marginal fitness distribution $\mathcal{N}^a(f)$:

$$\mathcal{N}^a(f) = 2(1-f) + \delta\Big\{ -4(1-f)\sum_{k=1}^{+\infty}\frac{1}{k^4\pi^4}\cos(k\pi f) + \\ + 4\sum_{k=1}^{+\infty}\frac{1}{k^5\pi^5}\sin(k\pi f) - 4\sum_{k=1}^{+\infty}\frac{1}{k^5\pi^5}\cos(k\pi f)\sin(k\pi f)\Big\} + O(\delta^2). \quad (21)$$

Using the properties of the trigonometric functions, the sums appearing in Eq. 21 can be evaluated in terms of Bernoulli polynomials (Gradshteyn and Ryzhik 2014). Finally, we obtain:

$$\mathcal{N}^a(f) = \underbrace{2(1-f)}_{\mathcal{N}^0(f)} + \delta\underbrace{\left\{\frac{4}{3}(1-f)B_4(\frac{f}{2}) - \frac{8}{15}B_5(\frac{f}{2}) + \frac{4}{15}B_5(f)\right\}}_{\mathcal{N}^{\,\mathrm{I}}(f)} + O\left(\delta^2\right), \quad (22)$$

where $B_k(z)$ is the $k^{\mathrm{th}}$-order Bernoulli polynomial of the variable $z$. Equation 22 is composed of the purely neutral contribution $\mathcal{N}^a_0(f)$, and of the first order correction $\delta\mathcal{N}^a_{\mathrm{I}}(f)$ with respect to $\delta$. Computing the average fitness of the population over the perturbative solution, we then obtain the linear expression:

$$\phi = \frac{1}{3} + \frac{1}{189}\,\delta + O\left(\delta^2\right), \quad (23)$$

as reported in the main text. Equation 22 can be used to estimate the threshold value $\delta$ at which the intermediate maximum appears in the marginal fitness distribution. The first derivative of Eq. 22 reads:

$$\frac{\partial\mathcal{N}^a(f)}{\partial f} = -2 + \frac{1}{90}\delta\Big\{75f^4 - 60f^3 - 90f^2 + 60f + 4\Big\}. \quad (24)$$

A local maximum exists whenever $\frac{\partial\mathcal{N}^a(f)}{\partial f} = 0$, that is whenever the following condition is satisfied:

$$75f^4 - 60f^3 - 90f^2 + 60f + 4 = \frac{180}{\delta}, \quad (25)$$

which is treated by graphical comparison in Fig. 1. The minimum value of $\delta$ for which the lines $y = \frac{180}{\delta}$ and $y = 75f^4 - 60f^3 - 90f^2 + 60f$ intersect, correspond to the local maximum of the latter.

The local maximum is located at $x = \frac{\sqrt{6}-1}{5}$ and has value of $\simeq 12.9$. Hence, the threshold value $\delta_{\text{th}}$ predicted by the perturbative approximation is:

$$\delta_{\text{th}} = \frac{180}{12.9} \simeq 13.95. \tag{26}$$

Below $\delta_{\text{th}}$, there is no solution to Eq. 25, and the marginal fitness distribution $\mathcal{N}^a(f)$ is monotonically decreasing (*redundancy-dominated* regime). Above $\delta_{\text{th}}$, the condition Eq. 25 has two solutions, indicating the presence of a new pair of local maximum and mininum. This estimation is consistent with the diagram of extrema positions obtained using the exact spectral solution (Fig. 4 of main text). The perturbative solution also gives a good estimation of the average fitness $\phi$, up to $\delta \simeq 30$ (left panel Fig. 2), that is when the linear prediction given by Eq. 23 is accurate. For large values of $\delta$, second order contributions due to selection become important and the perturbative solution fails. Indeed, in the right panel of Fig. 2 we can see that spectral (solid lines) and perturbative (dashed lines) profiles of the marginal fitness distributions agree almost perfectly for small $\delta$ (black line $\delta = 10$ case), and are qualitative consistent for intermediate values (black line $\delta = 30$ case). However, for higher values the approximation breaks down and also stops having biological value, as the distribution attains forbidden negative values (blue line $\delta = 50$ case).

#### Marginal fitness dynamics

Here we show how Eq. 11 of the main text, describing the dynamics of the marginal fitness distribution $\mathcal{N}^a(f)$ in the case of asymmetric landscapes, is obtained. For a general boundary profile $\mathcal{B}(x)$, integrating the RME over the neutral variable $y$, we obtain:

$$\int_0^{\mathcal{B}(x)} \frac{d\, n(x,y;t)}{dt}\, dy = \dot{\mathcal{N}}^a(x;t) = \mu \int_0^{\mathcal{B}(x)} \frac{\partial^2\, n(x,y;t)}{\partial\, x^2}\, dy$$

$$+ \mu \int_0^{\mathcal{B}(x)} \frac{\partial^2\, n(x,y;t)}{\partial\, y^2}\, dy + \gamma \mathcal{N}^a(x;t)\Big(F(x) - \overline{F}[\mathcal{N}^a(x;t)]\Big). \tag{27}$$

The second integral term on the right hand side of Eq. 27 is straightforward to handle. Applying the Fundamental Theorem of Integral Calculus to the second integral term, we obtain:

$$\int_0^{\mathcal{B}(x)} \frac{\partial^2\, n(x,y;t)}{\partial\, y^2}\, dy = \frac{\partial\, n(x,y;t)}{\partial\, y}\, \Big|_{y=0}^{y=\mathcal{B}(x)} = \frac{\partial\, n(x,y;t)}{\partial\, y}\, \Big|_{y=\mathcal{B}(x)}, \tag{28}$$

where we have used the no-flux boundary condition Eq. 10 in the last passage.

Performing a similar calculation on the first integral term of the right hand side of Eq. 27 and using the no-flux boundary condition Eq. 10, we obtain the following equality:

$$\int_0^{\mathcal{B}(x)} \frac{\partial^2\, n(x,y;t)}{\partial\, y^2}\, dx = \frac{d^2\, \mathcal{N}^a(x;t)}{dx^2} - 2\mathcal{B}'(x)\frac{\partial\, n(x,y;t)}{\partial x}\Big|_{y=\mathcal{B}(x)} + \mathcal{B}'^2(x)\frac{\partial n(x,y;t)}{\partial y}\Big|_{y=\mathcal{B}(x)} - \mathcal{B}''(x) n(x,y;t)_{y=\mathcal{B}(x)} \tag{29}$$

Finally, substituting Eqs. 28-29 into Eq. 27, we obtain Eq. 11 of the main text (recall that in the asymmetric landscape $f = x$).

### D Trait distribution on symmetric redundant landscapes

#### Spectral solution

In the symmetric case, the dynamics takes place in the phenotype space $\mathcal{P}_2 = \{x, y \mid x^2 + y^2 \leq 1\}$. The optimal phenotype occupies the centre $(0,0)$, and has maximum fitness value $f = 1$. Fitness decreases proportionally to the distance from the optimum. Hence, it is useful to describe the dynamics in the polar coordinates $(r, \theta)$, obtained from the trait variables via the usual coordinates transformation $x = r\,\cos\theta$, $y = r\,\sin\theta$. Describing the problem in polar coordinates $(r, \theta)$, we recognise $r$ as the selective variable, and $\theta$ as the neutral one: phenotypes lying on the same circle (i.e. with same $r$) will share the same fitness value, regardless of their value of $\theta$, as shown in Fig. 1, panel **b** the main text.

Denoting with $\nabla^2$ the Laplacian operator in polar coordinates, the stationary solution $n(r, \theta)$ of the RME in polar coordinates reads:

$$\mu\,\nabla^2\,n\,(r,\theta)\;+\;\gamma\,(F(r)-\phi)\,n\,(r,\theta) = \tag{30}$$

$$\mu\left\{\frac{\partial^2}{\partial r^2}+\frac{1}{r}\frac{\partial}{\partial r}\;+\;\frac{1}{r^2}\frac{\partial^2}{\partial\theta^2}\right\}n(r,\theta)\;+\;+\gamma\,(F(r)-\phi)\,n\,(r,\theta)=0, \tag{31}$$

and subject to the usual conditions:

$$\begin{cases} \int_0^1\int_0^{2\pi}\;n(r,\theta)\,r\,drd\theta = 1 & \textit{population normalisation} \\ \frac{\partial\,n(r,\theta)}{\partial\,r}\;|_{r=1}=0 & \textit{no-flux} \\ \phi=\int_0^1\int_0^{2\pi}\;F(r)\,n(r,\theta)\,r\,drd\theta & \textit{self-consistent average fitness.} \end{cases} \tag{32}$$

Differently from the asymmetric case, separation of variables can be applied since the *no-flux* boundary condition only depends on the selective variable $r$. Hence, the solution can be found in the form $n(r, \theta) = R(r)Y(\theta)$, where radial $R(r)$ and angular $Y(\theta)$ contribution are factorised and describe, respectively, selective and neutral contributions to the phenotype distribution. Moreover, the angular contribution is trivially constant, due to circular symmetry, and can then be absorbed in the constant of normalisation. Hence, we are left with the radial equation for the selective contribution:

$$\frac{\partial^2 R(r)}{\partial r^2}+\frac{1}{r}\frac{\partial R(r)}{\partial r}+\delta\,R(r)\,(F(r)-\phi)=0 \tag{33}$$

with

$$\begin{cases} \int_0^1 R(r)\, r\, dr = 1 \\ \frac{\partial R(r)}{\partial r}\mid_{r=1} = 0 \\ \phi = \int_0^1 F(r)\, R(r)\, r\, dr, \end{cases} \tag{34}$$

Similarly to the previous problem, we look for a solution in the form of a linear combination of the eigenfunctions of Helmholtz problem on the disk $\mathcal{G}_{\mathrm{Sym}}$ with Neuman boundary conditions:

$$R(r) = \sum_k c_k\, J_0\left(\beta_{0,k}\, r\right), \tag{35}$$

where $J_0$ is the Bessel function of first kind for order zero, and $\beta_{0,k}$ is the positive $k^{\mathrm{th}}$ root of the first derivative $J_0'$ (Grebenkov and Nguyen 2013). Again, it is straightforward to show that the first coefficient $c_0 = 2$ is fixed by the normalisation constraint. Substituting Eq. 35 into Eq. 33 and exploiting the orthonormalisation properties of the eigenfunctions (Bowman 2012), we obtain the linear system of equations for the other coefficients $\{c_k\}$ of the expansion:

$$c_l\left(-\delta\,\phi - \beta_l^2\right)\frac{J_0(\beta_l)^2}{2} + \delta\sum_{k=1}^{+\infty} c_k\,\langle\, F(r)\,\rangle_{k,l} = 0, \tag{36}$$

with $\langle\, F(r)\,\rangle_{k,l} = \int_0^1 F(r)\, J_0\left(\beta_k r\right)\, J_0\left(\beta_l r\right)\, r\, dr$. Similarly to the procedure employed for the asymmetric case, truncating the system of equations to a finite number of terms, one first solves the above linear problem with standard numerical routines, then finds $\phi$ solving the self-consistent condition on the average fitness.

### Marginal fitness dynamics

In the symmetric case, the marginal fitness distribution can be straightforwardly found from the radial contribution $R(r)$, as integrating over the neutral variable $\theta$ one obtains:

$$\mathcal{N}^s(r) = \int_0^{2\pi} n(r,\theta)\, r\, d\theta = \int_0^{2\pi} R(r) Y(\theta)\, r\, d\theta = r\, R(r). \tag{37}$$

Hence, its dynamics can directly be derived by inserting first and second derivates of expression Eq. 37 into the radial Eq. 33, and we obtain:

$$\frac{\partial^2 \mathcal{N}^s(r)}{\partial r^2} + \frac{\partial}{\partial r}\left[v(r)\mathcal{N}^s(r)\right] + \delta\,\mathcal{N}^s(r)\,(F(r)-\phi) = 0, \qquad \text{with} \quad v(r) = -\frac{1}{r}. \tag{38}$$

Recalling that the selective variable is $f = 1 - r$, one then obtains Eq. 9 of the main text.

#### Generalisation to higher dimensions

The radial Eq. 33 describing the dynamics in the two-dimensional symmetric landscape can be easily generalised to higher dimensions, where again selection is encoded in the radial coordinate, while the remaining $D-1$ angular coordinates are neutral. In dimension $D$, the radial equation for the stationary solution of the RME will be:

$$\frac{\partial^2 R(r)}{\partial r^2} + \frac{D-1}{r}\frac{\partial R(r)}{\partial r} + \delta\,(F(r)-\phi)\,R(r) = 0, \tag{39}$$

with boundary conditions:

$$\begin{cases} \int_0^1 r^{D-1}\,R(r)\,dr = 1 \\ \frac{\partial R(r)}{\partial r}\,|_{r=1} = 0 \\ \phi = \int_0^1 F(r)\,R(r)\,r^{D-1} dr. \end{cases} \tag{40}$$

Then, the corresponding marginal fitness distribution will read:

$$\mathcal{N}_D(r) = r^{D-1}\,R(r). \tag{41}$$

In general, to solve Eq. 39 one can employ the same machinery we have discussed throughout this work: expand in the eigenfunctions of the generalised Laplacian in dimension $D$; truncate to a finite order the system on linear equations for the coefficients of the expansion; find the proper $\phi$ with the self-consistent condition.

Interestingly, for $D = 3$ (that is for a spherical adaptive space) and in the case of linear fitness function $F(r) = 1 - r$, Eq. 39 can be solved directly. Performing the change of variable $\tilde{R}(r) = rR(r)$, the radial equation becomes an Airy equation (similar to Eq. 4):

$$\frac{\partial^2 \tilde{R}(r)}{\partial r^2} + \delta\,(\phi - r)\,\tilde{R}(r) = 0, \tag{42}$$

and the solution is again a linear combination of the Airy functions. Finally, according to Eq. 41 the marginal fitness distribution on the sphere will be:

$$\mathcal{N}(r) = \mathcal{Z}\, r \left\{ \mathrm{Ai}\left([r - \phi]\sqrt[3]{\delta}\right) + \mathcal{C}\,\mathrm{Bi}\left([r - \phi]\sqrt[3]{\delta}\right) \right\}, \quad \delta = \frac{\gamma}{\mu} \tag{43}$$

with $\mathcal{Z}, \mathcal{C}$ normalisation and no-flux boundary conditions.

### E Numerical Simulations

Throughout this work, the theoretical fitness distributions predicted by the stationary solutions of the RME have been compared with numerical agent-based simulations, where mutation and competition events are modelled by birth-death stochastic processes preserving the total population number. The scheme we employed is based on a standard Gillespie algorithm. The algorithm can be applied to simulate any mutation-selection dynamics with continuous variables, regardless of the choice of the fitness function, of the dimension and shape of the adaptive space. The codes used in this work are freely available here

The population is composed of $N$ individuals, each individual is labelled with a phenotype $\underline{\mathbf{p}}$ whose components are continuous variables $\in [0, 1]$. A time step is composed of $N$ events of either mutation or competition. At rate $\hat{\mu}$, an individual of the population is selected at random and its phenotype $\underline{\mathbf{p}}$ undergoes a mutation reaction:

$$\underline{\mathbf{p}} \xrightarrow{\hat{\mu}} \underline{\mathbf{p}} + \underline{\zeta}, \tag{44}$$

where $\underline{\zeta}$ is a vector of random variables extracted from a probability distribution $\mathcal{P}(0, \sigma^2)$ with *zero* mean and variance $\sigma^2$. If the new mutant phenotype would cross the border of the adaptive space, the mutation is rejected and the individual retains its original phenotype, that is reflecting boundary conditions are considered. The assumption of *zero* mean implies that mutations are unbiased, that is the adaptive space is explored with same probability in any direction, and no distinction between deleterious, beneficial or neutral mutations is made. For the analysis provided in this work we have used a uniform probability distribution, but we have checked that the results hold for a Gaussian distribution as well. For small values of $\sigma^2$, the accumulation of such mutations leads the individuals to perform a random walk exploration of the adaptive space (see e.g. Wakano et al. 2017 for details). Thus, for large population mutations are be captured by the local diffusion operator $\nabla^2$, with effective mutation rate given by the diffusion coefficient $\mu = \frac{\sigma^2}{2}\hat{\mu}$, and no-flux boundary conditions.

Concurrent with the above mutation process, at rate $\hat{\gamma}$, two individuals with phenotype $\underline{p}_1$ and $\underline{p}_2$ are chosen at random and undergo a competition event: one individual perishes and the other reproduces, with probability proportional to the difference between their fitness. The two possible outcomes are schematised by the complementary pair of reactions:

$$\underline{p}_1 + \underline{p}_2 \xrightarrow{\hat{\gamma}\left(\frac{1}{2} + \omega\frac{F(\underline{p}_1) - F(\underline{p}_2)}{2}\right)} 2\,\underline{p}_1$$

$$\tag{45}$$

$$\underline{p}_1 + \underline{p}_2 \xrightarrow{\hat{\gamma}\left(\frac{1}{2} + \omega\frac{F(\underline{p}_2) - F(\underline{p}_1)}{2}\right)} 2\,\underline{p}_2$$

The reaction rates in Eq. 45 are composed of two probabilities: a baseline probability of $\frac{1}{2}$ equal for both, and a term proportional to the fitness difference. The constant of proportionality $\omega \in [0, 1]$ is called *selection strength*, and can be used to modulate the importance of fitness during the competition for reproduction. For $\omega = 0$, the dependence on fitness disappears and the two reactions occur with same probability: hence, $\omega = 0$ describes a *purely neutral* scenario. For $\omega \neq 0$, the outcome of competition is skewed in favour of the fitter individual.

This formulation of selection relies only on the *local* information of the individuals involved and does not depend on macroscopic quantities such as the average fitness $\bar{f}$ of the population. In game theory, interpreting fitness as payoffs, the above formulation is sometimes called *local update rule* (Traulsen et al. 2005). For large populations, iteration of the selection event leads to the nonlocal selection term appearing in Eq. 3 of the main text, which is also known as *replicator term*, with effective competition rate $\gamma = 2\,\omega\,\hat{\gamma}$.

### References


Abramowitz, M. and I.A. Stegun (1964). *Handbook of mathematical functions: with formulas, graphs, and mathematical tables.* Vol. 55. Courier Corporation.

Bowman, Frank (2012). *Introduction to Bessel functions.* Courier Corporation.

Damle, Anil and Geoffrey Colin Peterson (2010). "Understanding the eigenstructure of various triangles". In: *SIAM Undergraduate Research Online* 3.1, pp. 187–208.

Gradshteyn, Izrail Solomonovich and Iosif Moiseevich Ryzhik (2014). *Table of integrals, series, and products.* Academic press.

Grebenkov, Denis S and B-T Nguyen (2013). "Geometrical structure of Laplacian eigenfunctions". In: *siam REVIEW* 55.4, pp. 601–667.

Traulsen, Arne, Jens Christian Claussen, and Christoph Hauert (2005). "Coevolutionary dynamics: from finite to infinite populations". In: *Physical review letters* 95.23, p. 238701.

Wakano, Joe Yuichiro, Tadahisa Funaki, and Satoshi Yokoyama (2017). "Derivation of replicator–mutator equations from a model in population genetics". In: *Japan Journal of Industrial and Applied Mathematics* 34.2, pp. 473–488.

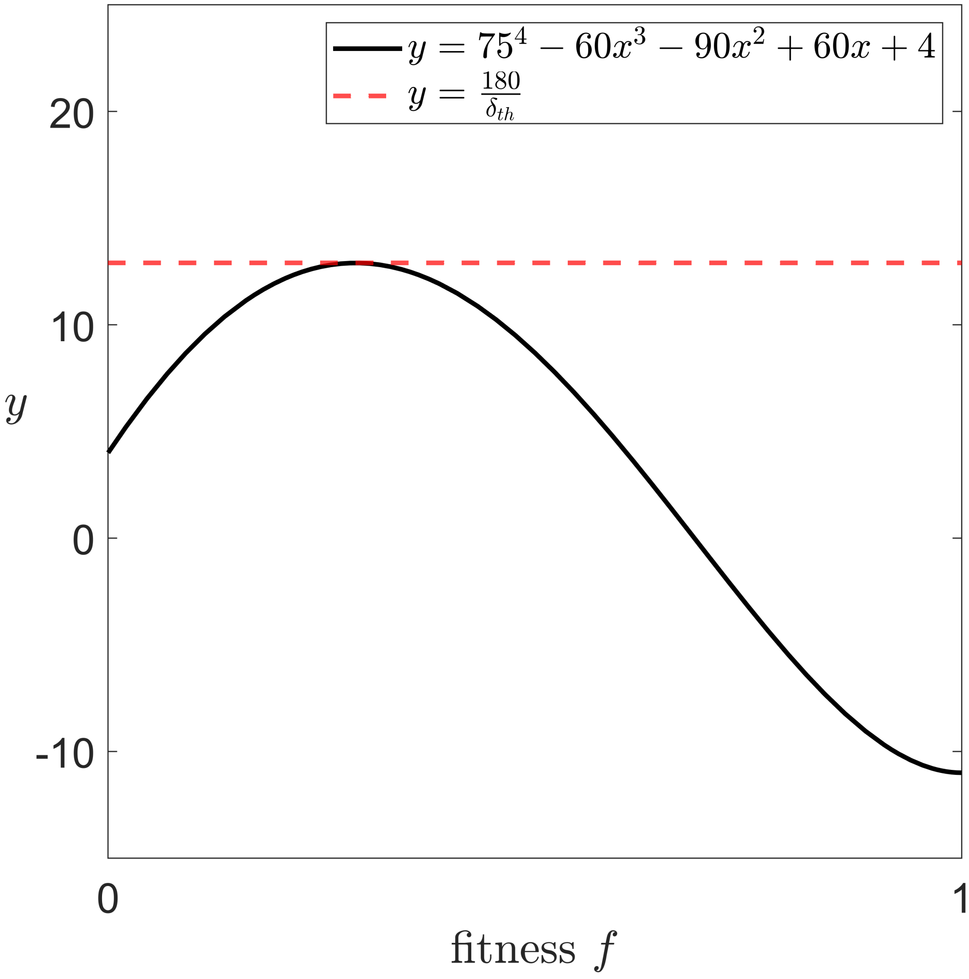


Supplementary Figure 1: **Perturbative estimation of the threshold value $\delta_{\mathbf{th}}$.** The intersection between the left-hand side (black line) of Eq. 25 and horizontal lines gives the position of the local maxima and minima of the marginal fitness profiles, under the perturbative approach. The value of $\delta$ for which they intersect for the first time corresponds to the threshold value $\delta_{\text{th}}$ (red dashed line) predicted by the perturbative approach. Below $\delta_{\text{th}}$, there is no intersection, hence no local extrema are present in the interior of the genomic space. Above $\delta_{\text{th}}$, horizontal lines intersect twice the black line, identifying a pair of local maxima and minima, in agreement with the behaviour of the spectral solutions.

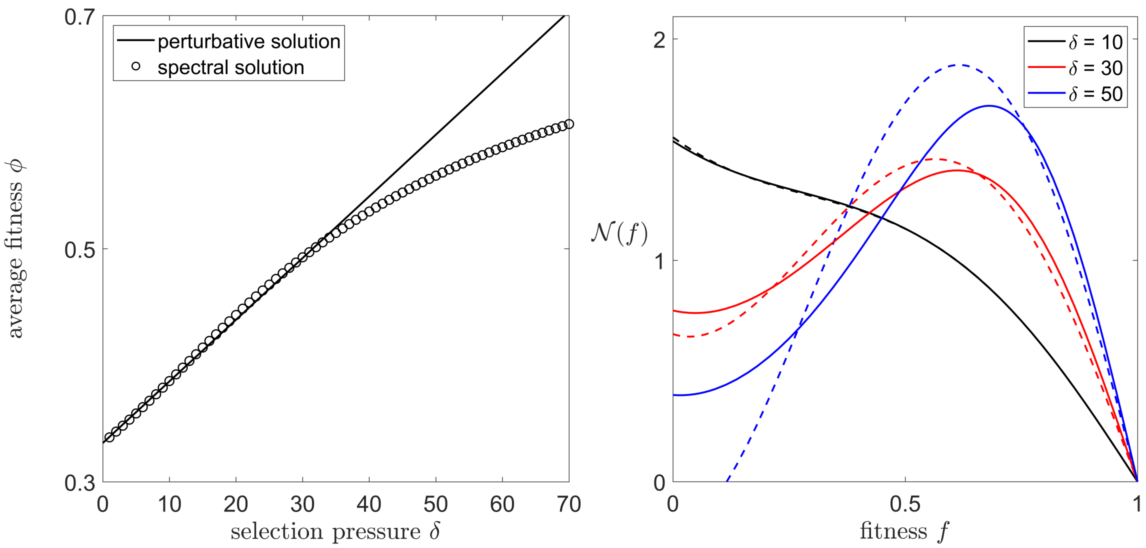


Supplementary Figure 2: **Perturbative vs spectral predictions.** Left panel: Comparison between perturbative (black line) and spectral (circles) estimation of the average fitness $\phi$, at the varying of $\delta$, in the case of asymmetric landscape. The perturbative solution gives a good estimate up to $\delta \simeq 30$. Right panel: Comparison between perturbative (dashed lines) and spectral (solid lines) profiles of the marginal fitness distribution. As we can see, the perturbative solution works very well for small $\delta$ (black lines), that is in the *redundancy-dominated* regime. For intermediate values of $\delta$ (red lines), the two profiles start diverging, yet the perturbative one still captures the qualitative behaviour. For large values of $\delta$ (blue lines), the perturbative approximation breaks down, giving negative non-biological profiles.
